# A lipid nanoparticle-mRNA vaccine provides potent immunogenicity and protection against *Mycobacterium tuberculosis*

**DOI:** 10.1101/2024.07.31.605765

**Authors:** Hannah Lukeman, Hareth Al-Wassiti, Stewart A. Fabb, Leonard Lim, Trixie Wang, Warwick J. Britton, Megan Steain, Colin W. Pouton, James A. Triccas, Claudio Counoupas

## Abstract

*Mycobacterium tuberculosis* remains the largest infectious cause of mortality worldwide, even with over a century of widespread administration of the only licensed tuberculosis (TB) vaccine, Bacillus Calmette-Guérin (BCG). mRNA technology remains an underexplored approach for combating chronic bacterial infections such as TB. We have developed a lipid nanoparticle (LNP)-mRNA vaccine encoding for a fusion protein of two immunogenic TB antigens, termed mRNA^CV2^. In C57BL/6 mice intramuscularly vaccinated with mRNA^CV2^, high frequencies of polyfunctional, antigen-specific Th1 CD4^+^ T cells were observed in the blood and lungs, which was associated with the rapid recruitment of both innate and adaptive immune cells to lymph nodes draining the site of immunisation. mRNA^CV2^ vaccination provided significant pulmonary protection in *M. tuberculosis*-infected mice, reducing bacterial load and inflammatory infiltration in the lungs. As BCG is widely administered in infants in TB endemic countries, new TB vaccines should be able to boost the effects of BCG. Importantly, mRNA^CV2^ enhanced immune responses and long-term protection when used to boost BCG-primed mice. These findings, which provide the first report of a highly protective LNP-mRNA vaccine for TB, highlight the potential of the LNP-mRNA platform for TB control and support further research to facilitate translation to humans.

## Introduction

Despite the existence of curative antibiotics and a licensed vaccine, *Mycobacterium tuberculosis* remains a major burden on global health. In 2022, tuberculosis (TB) reclaimed its position as the leading cause of death by a single infectious agent, with an estimated 10.6 million new cases and 1.3 million deaths^1^. Furthermore, approximately a quarter of the global population is latently infected with *M. tuberculosis*, representing a large reservoir of individuals who may reactivate TB later in life^1,2^. The ongoing global health emergency caused by TB is partly due to the variable efficacy of the *Mycobacterium bovis* bacillus Calmette-Guérin (BCG) vaccine. Whilst generally protective in children, efficacy of BCG ranges between 0-80% against pulmonary manifestations of the disease in adults^3,4^. Human immunodeficiency virus (HIV) coinfection, growing rates of drug resistance, and a lack of adherence or access to treatments further hinders control of TB^5^. Development of a more efficacious vaccine is critical for overcoming these challenges and reducing the global TB burden^6^.

A number of TB vaccine candidates are currently in clinical trials. They comprise of a wide range of delivery platforms, notably protein subunit, live attenuated, inactivated, and viral vectored-vaccines^7^. The only clinically protective candidate to date is the protein subunit vaccine M72/AS01_E_, which demonstrated in a Phase 2 trial an efficacy of 49.7% for preventing progression to active disease in adults latently infected with *M. tuberculosis*^8^. An ongoing Phase 3 trial will determine if this vaccine can achieve the World Health Organisation (WHO) preferred product characteristics for a new TB vaccine, including at least 50% protection in subjects with and without evidence of latent *M. tuberculosis* infection, in different geographical regions^9^. As such, further research into more efficacious TB vaccines is required. Particularly exploring novel approaches, given the current limitations of existing vaccines and the urgent need for enhanced protection and long-term immunity against *M. tuberculosis*.

The successes of the lipid nanoparticle (LNP)-mRNA platform following the SARS-CoV-2 pandemic supports investigating their utility against TB, and WHO has identified this platform as a crucial tool to potentially reduce the global burden of TB^6,10^. Unlike protein subunit or live vaccines, mRNA vaccines are produced in a cell-free environment^11^. This allows for rapid, scalable, and cost-effective development, rendering them suitable for widespread administration in low-income countries that have the greatest burden of *M. tuberculosis*^12^. Furthermore, mRNA vaccines are non-infectious, permitting use in immunocompromised individuals (i.e. HIV-infected) for whom live vaccines may be problematic^13^. mRNA vaccine technology has proven effective in generating robust immune responses against viral pathogens, such as in the case of SARS-CoV-2, where humoral immunity is the key correlate of protection^14^. However, its application against chronic bacterial infections such as TB remains uncertain. This is because effective TB immunity requires the stimulation of multifunctional CD4^+^ T cells to limit bacterial growth and persistence, presenting a unique challenge against a pathogen that is highly evolved to evade immune detection and clearance^15,16^.

In the present study, we developed an mRNA vaccine encoding for CysVac2, termed mRNA^CV2^. CysVac2 is a fusion protein of two *M. tuberculosis* antigens; the secreted immunodominant Ag85B, and CysD, a component of the sulphur assimilation pathway that is overexpressed in chronic stages of infection^17^. CysVac2 is immunogenic and protective in mice when used in adjuvanted subunit vaccine formulations against *M. tuberculosis*^17–20^. Here, we demonstrate that the mRNA^CV2^ vaccine induces potent antigen-specific Th1-biased immune responses and is protective against *M. tuberculosis* in a murine model, both alone and as a booster vaccination to BCG. This study demonstrates the potential of mRNA vaccine technology in combating chronic infections such as TB, offering a promising avenue for future vaccine development.

## Methods

### Bacterial strains

*M. tuberculosis* H37Rv and BCG Pasteur were cultured at 37 °C in Middlebrook 7H9 medium (BD) supplemented with 0.5% glycerol, 0.02% Tyloxapol and 10% albumin-dextrose-catalase (ADC), or solid Middlebrook 7H10 medium (BD) supplemented with oleic acid-ADC, respectively.

### LNP-mRNA production

mRNA was produced using the HiScribe T7 mRNA synthesis kit (NEB, Australia) using a template of linearized DNA produced by PCR amplification. Transcribed mRNAs included 3’-UTR with Kozak sequence, 5’-UTR, and polyA_125_ tails. The sequences were codon-optimized to reduce the uridine content. In addition, N1-methyl-pseudoUTP was used instead of UTP to produce chemically modified mRNA, in common with the strategy used for the two approved COVID-19 vaccines. CleanCap reagent AG (TriLink) was used in accordance with the manufacturer’s recommendations to produce Cap1 chemistry at the 5’ terminus. The mRNA was subject to cellulose purification before use.

The following lipids were used to produce LNPs; the ionizable lipid (6*Z*,9*Z*,28*Z*,31*Z*)-Heptatriaconta-6,9,28,31-tetraen-19-yl-4-(dimethylamino)butanoate (‘DLin-MC3-DMA’, MedChemExpress, USA), cholesterol (Sigma-Aldrich, Germany), 1,2-distearoyl-sn-glycero-3-phosphocholine (DSPC) (Avanti Polar Lipids Inc., USA), and 1,2-dimyristoyl-rac-glycero-3-methoxypolyethylene glycol-2000 (DMG-PEG2000) (Avanti Polar Lipids Inc., USA) (Supplementary Fig. 1). The molar percentages of lipids used in the formulation were as follows: DLin-MC3-DMA: cholesterol: DSPC: DMG-PEG2000 (50: 39.85: 10: 0.15). Manufacture of LNPs involved the following steps: an aqueous solution of mRNA at pH4 was mixed with a solution of the four lipids in ethanol, using a microfluidics mixing device (NxGen Ignite Nanoassemblr, supplied by Precision Nanosystems, Vancouver, Canada). The suspension of nanoparticles was adjusted to pH 7.4 using a 1:3 dilution in Tris buffer, then dialysed against 25mM Tris buffer to remove the residual ethanol. The LNP suspension was adjusted with sucrose solution to produce the cryoprotected, isotonic final form of the product. The product was filtered (0.22µm) prior to being aliquoted into sterile vials for storage at -80°C. Characterization of the LNPs included analysis for RNA content, encapsulation efficiency, and RNA integrity. Particle size and polydispersity index (PDI) were determined by dynamic light scattering using a Zetasizer (Malvern Instruments). Typically, encapsulation efficiency was 85-95%, particle size (Z-average) of the 0.15% PEGylated lipid formulation was 120-160nm with polydispersity index < 0.2

### Mice and immunisation

Female C57BL/6 mice (6-8 weeks of age) were purchased from Australian BioResources (Mossvale, Australia) and housed under specific pathogen-free conditions at the Centenary Institute Bioresources animal facility (Sydney, Australia) under Biosafety Level (BSL) II or III conditions. All experiments were performed with approval of the Sydney Local Health District Animal Ethics and Welfare Committee (protocol 2020-009). For all vaccinations, mice were anaesthetised with gaseous isoflurane (3-4%, O_2_ 1 L/min). For early immune kinetics experiments, mice were vaccinated once intramuscularly using insulin syringes with endotoxin-free PBS (50 μL each hind leg; Sigma-Aldrich) or mRNA^CV2^ (5 μg total, 50 μL volume each hind leg). For standard immunogenicity and protection experiments, mice were vaccinated once subcutaneously with 5 × 10^5^ colony forming units (CFU) of BCG (200 μL in endotoxin-free PBS), or intramuscularly 3 times at 3-week intervals in alternating hind legs with endotoxin-free PBS (50 μL) or mRNA^CV2^ (5 μg, 50 μL total volume, single leg). For BCG boosting experiments, mice were either vaccinated with BCG as above, or left unvaccinated, then 6-7 months later selected mice were vaccinated intramuscularly with two doses of mRNA^CV2^ (5 μg, 50 μL total volume, alternate single leg), 3 weeks apart.

### M. tuberculosis infection, bacterial quantification and histology

Five weeks after final vaccinations, mice were challenged with *M. tuberculosis* H37Rv (infective dose of ∼100 viable bacilli per mouse) via aerosol using a Middlebrook airborne infection apparatus (Glas-Col). After 4 or 21 weeks, the left lung and spleen were harvested, homogenised and serially diluted in PBS, before plating on supplemented Middlebrook 7H10 agar plates. These were incubated at 37 C and 5% CO_2_ for 3-4 weeks before CFU were enumerated and expressed as either Log_10_ CFU or Log_10_ CFU ± difference of each individual mouse compared to the mean of unvaccinated mice.

For histological examination, the middle right lobe from infected mice was perfused with 10% buffered formalin, embedded in paraffin, sectioned, and stained with haematoxylin and eosin. Slides were observed with a LeicaDM microscope (Leica Microsystems, North Ryde, Australia) at 10× and 40× magnification, and acquired as a mosaic.

### Sample collection and processing

Blood was collected from the lateral tail vein into heparin (20 U, Sigma-Aldrich). Plasma was separated by centrifugation and peripheral blood mononuclear cells (PBMCs) were isolated by stratification on Histopaque 1083 (Sigma-Aldrich). Following euthanasia by CO_2_ asphyxiation, organs were collected into RPMI 1640 (Life Technologies, ThermoFisher Scientific, Sydney, Australia). Lungs were perfused with PBS prior to collection and mechanically dissociated with the GentleMACS dissociator (Miltenyi Biotec, Sydney, Australia) before enzymatic digestion with DNAse I (10 U/mL; Sigma-Aldrich) and Collagenase IV (10 U/mL; Sigma-Aldrich) for 45 min at 37 °C. Single-cell suspensions were prepared by filtering through a 70 μm nylon cell strainer and red blood cells were lysed with ACK lysis buffer (ThermoFisher Scientific) before washing and resuspension in RPMI 1640 (Life Technologies, ThermoFisher Scientific) supplemented with 10% FCS (Sigma), 2-ME (0.05 mM), and penicillin-streptomycin (100 U/mL; Sigma), henceforth referred to as cRPMI. Lymph nodes undergoing myeloid cell flow cytometric staining were initially enzymatically digested with DNAse I (10 U/mL; Sigma-Aldrich) and Collagenase IV (10 U/mL; Sigma-Aldrich) for 30 min at 37 °C. Otherwise, lymph nodes were directly filtered through a 70 μm nylon sieve, centrifuged to pellet cells, and resuspended in cRPMI.

For restimulation of antigen-specific cells, the CysVac2 fusion protein was produced by recombinant expression in *E. coli* as previously described^17^. Cells were incubated with 5-10 μg/mL CysVac2 in cRPMI for 4 hrs at 37 °C and 5% CO_2_, followed by Protein Transport Inhibitor Cocktail (Life Technologies, Thermo Fisher Scientific) overnight at 37 °C, 5% CO_2_. Unstimulated cells were stained immediately after collection.

### Flow cytometry

PE-conjugated Ag85B_240-254_:I-A^b^ Class II MHC tetramer was provided by the NIH Tetramer Core Facility (Atlanta, USA). If undergoing tetramer staining, unstimulated single cell suspensions were initially incubated with tetramer diluted 1:150 in cRPMI containing Fc-block (1:200, anti-CD16/CD32; clone 2.4G2; Becton Dickinson) for 45 min at 37 °C, 5% CO_2_. When used, CCR7, XCR1 and CXCR3 staining was also performed at this step. Cell surface markers were stained with monoclonal antibodies (mAbs) outlined in Supplementary Table 1, Fixable Blue Dead Cell stain (1:300, Life Technologies, Thermo Fisher) and Fc-block diluted in FACS wash (PBS + 2% FCS + 5 mM EDTA) for 30 min at 4 °C. After washing, if intracellular cytokine staining was being performed, cells were permeabilised with BD Cytofix/Cytoperm Fixation/Permeabilization Kit (Becton Dickinson) as per the manufacturer’s instructions; or, if undergoing staining for transcription factors, with eBioscience^TM^ Foxp3/Transcription Factor Staining Buffer Set (Invitrogen) as per the manufacturer’s instructions. This was followed by staining with mAbs for intracellular markers for 30 min at 4° C. Cells were. acquired on LSRII 5L cytometer (BD biosciences, Franklin Lakes, NJ, USA) and analysed using FlowJo^TM^ analysis software (Treestar, USA) using the gating strategies outlined in Figures S1-3. To quantify frequencies of single-, double-, and triple-positive cytokine-producing CD4^+^ T cell subsets, Boolean gating was performed.

### Antibody Enzyme-Linked Immunosorbent Assays (ELISAs)

96-well microplates (Corning, Sigma-Aldrich) were incubated overnight at room temperature (RT) with 0.5 μg/mL CysVac2 protein diluted in PBS. Free binding sites were blocked with 3% (w/v) BSA for 1-2 h at RT, plates were washed with 0.01% (v/v) Tween and 0.1% (w/v) BSA in PBS, and serially diluted serum samples were incubated for 45 min at 37 °C, 5% CO_2_. After washing, either biotinylated polyclonal goat anti-mouse IgG1 (1:50,000, Abcam) or goat anti-mouse IgG2c (1:10,000, Abcam) detection antibodies were incubated for 1 h at RT, followed by another wash and incubation with streptavidin-HRP (1:30,000, Abcam) for 30 min at RT. The reaction was developed in washed plates with 0.1 mg/mL 3,3’,5,5’-Tetramethylbenzidine (Sigma-Aldrich) and H_2_O_2_ in 0.05 M phosphate-citrate buffer, then stopped with 0.2 M H_2_SO_4_. Absorbances were read at 450 nm using the Tecan Infinite M1000 PRO plate reader. Endpoint titres were calculated in GraphPad Prism® version 10 (GraphPad Software Inc.) by fitting to a sigmoidal curve and determining the dilution of sample that reaches the mean absorbance of PBS-vaccinated control serum ± 3 SD.

### Statistical analysis

Statistical significance was calculated using GraphPad Prism® version 10 (GraphPad Software Inc., La Jolla, California). Unpaired t-tests or Mann-Whitney tests were utilised for comparisons between 2 groups. For multi-group comparisons, 1-way ANOVA or 2-way ANOVA were performed and corrected with Dunnett’s or Tukey’s *post-hoc* test where appropriate. A difference was determined as statistically significant when *p* ≤ 0.05.

## Results

### Vaccination with mRNA^CV2^ induces robust Th1-biased vaccine-specific cellular and humoral immunity

To determine if LNP-mRNA based delivery of TB antigens would be protective, a nucleoside-modified mRNA encoding for the CysVac2 fusion protein was constructed. This mRNA was encapsulated in a unique LNP formulation developed at Monash Institute of Pharmaceutical Sciences (MIPS). This formulation (see methods) has previously been used in a Phase 1 clinical trial of a SARS-CoV-2 mRNA vaccine^21^. To define the immune response elicited by this vaccine, termed mRNA^CV2^, C57BL/6 mice were vaccinated intramuscularly 3 times, at 3-week intervals with mRNA^CV2^ or vehicle, and antigen-specific production of cytokines by peripheral blood mononuclear cells (PBMCs) determined by flow cytometry (Supplementary Fig. 2), at one week post each vaccination (Fig. 1A). Subcutaneous immunisation with the BCG standard of care vaccine was used as a comparator.

**Figure 1:**
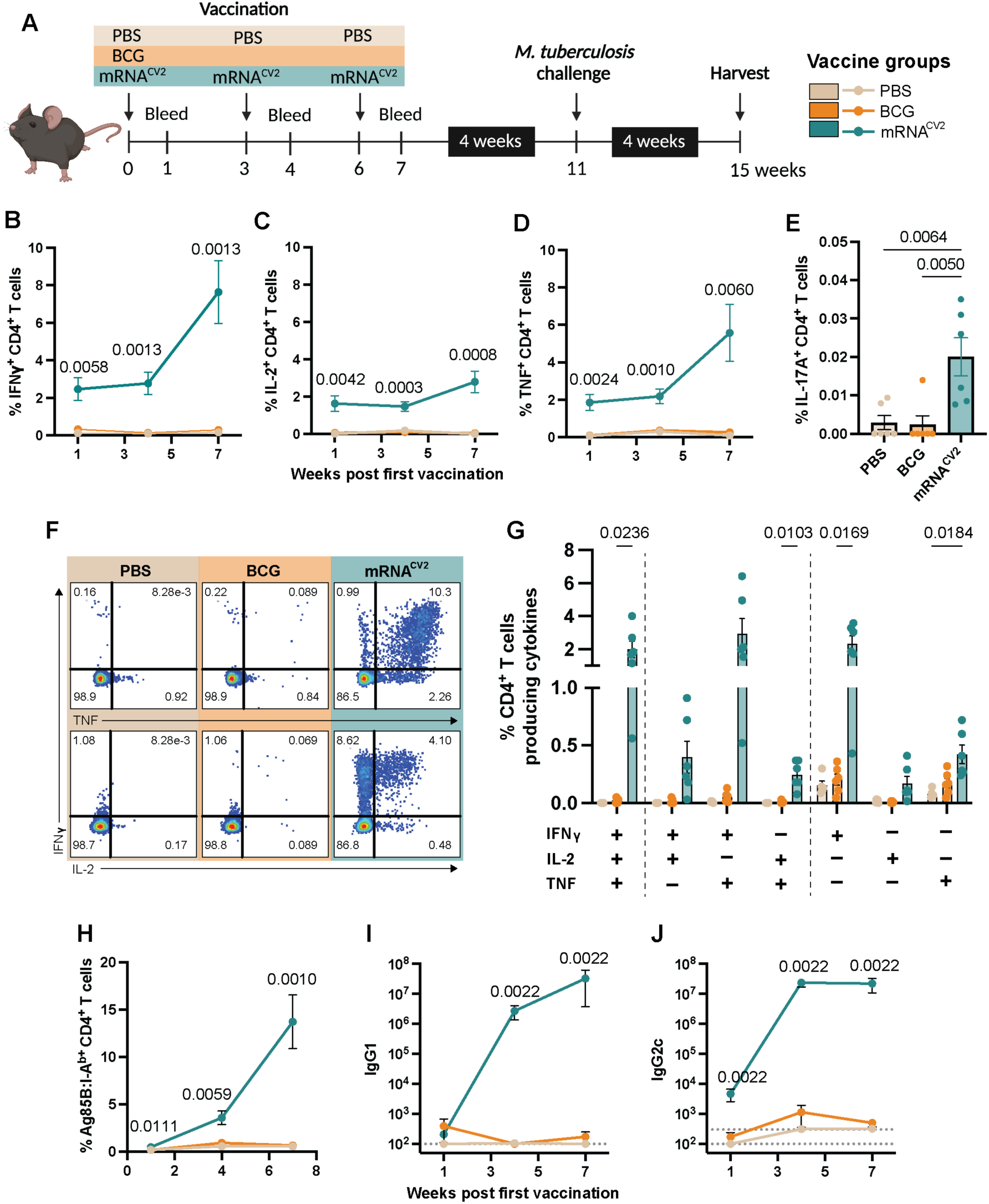
Potent circulating CysVac2-specific CD4^+^ T cell responses following mRNA^CV2^ vaccination. **(A)** Schematic of experimental outline. **(B-D)** Proportion of CysVac2-specific IFNγ-**(B),** IL-2-**(C)** or TNF-**(D)** producing CD4^+^ T cells in the peripheral blood of C57BL/6 mice a week after each vaccine dose, and **(E)** IL-17A-producing CD4^+^ T cells a week after the final vaccination. **(F)** Representative cytometric plots of CD4^+^ T cells producing cytokines. **(G)** Proportion of polyfunctional cytokine-producing CD4^+^ T cells 7 weeks after initial vaccination. **(H)** Proportion of Ag85B:I-A^b+^ CD4^+^ T cells a week after each vaccine dose. Endpoint titre of CysVac2-specific **(I)** IgG1 and **(J)** IgG2c antibodies in sera from mice a week after each vaccine dose. Dotted line is limit of detection. Data are expressed as mean ± SEM with 5-6 mice in each group and representative of 2 independent experiments. Significance between groups was determined by **(B-D, H)** multiple unpaired t-tests between BCG and mRNA^CV2^ vaccine groups, **(E)** One-way ANOVA corrected with Tukey’s *post-hoc* test, **(G)** Two-way ANOVA corrected with Tukey’s *post-hoc* test, and **(I, J)** Mann-Whitney test between BCG and mRNA^CV2^ vaccine groups.

The frequency of CD4^+^ T cells secreting characteristic Th1 cytokines in mice vaccinated with mRNA^CV2^ was significantly elevated above those in PBS- or BCG-vaccinated mice, with frequencies continuing to increase after each vaccination (Fig. 1b-d). IL-17A-producing CD4^+^ T cells were also significantly higher in the mRNA^CV2^ group, but at a much lower frequency compared to the other cytokines (Fig. 1e). Further analysis revealed a large proportion of triple- (IFNγ^+^, IL-2^+^, TNF^+^) or double-positive (IFNγ^+^, TNF^+^) populations induced by mRNA^CV2^ above that observed for PBS or BCG (Fig. 1f, g). Ag85B:I-A^b^ tetramer staining, specific for the Ag85B p25 epitope contained within mRNA^CV2^ ^17^, was employed to further define vaccine-specific CD4^+^ T cells in the blood (Supplementary Fig. 3). From a week after the first vaccination, there was a markedly enhanced population of Ag85B:I-A^b+^ CD4^+^ T cells in mice vaccinated with mRNA^CV2^ compared to PBS or BCG groups, the frequency of which increased with each subsequent dose to approximately 14% of CD4^+^ T cells (Fig. 1g). This demonstrates that mRNA^CV2^ induces strong, Th1-biased cellular immune response in vaccinated mice, with high levels of vaccine-specific cells circulating in the periphery.

We next examined if the strong T cell immunity induced by mRNA^CV2^ was accompanied by antibody responses, considering the emerging role of humoral immunity in protection against *M. tuberculosis* infection^19,22^. CysVac2-specific IgG antibodies were quantified in the plasma of vaccinated mice by ELISA. mRNA^CV2^ induced significantly greater production of both IgG1 and IgG2c than PBS or BCG, which was apparent after the 2^nd^ dose of vaccine (Fig. 1i, j). The titre of IgG2c peaked at an earlier timepoint than IgG1, with IgG1 requiring 3 rather than 2 doses of vaccine to reach an equivalent titre. Taken together, these data demonstrate that mRNA^CV2^ is highly immunogenic, capable of inducing robust cellular and humoral vaccine-specific responses in vaccinated mice, with a characteristic Th1-like phenotype.

### mRNA^CV2^ is protective in a mouse model of M. tuberculosis

Considering these strong immune responses following vaccination with mRNA^CV2^, we sought to determine if this was associated with protection against *M. tuberculosis*. Five weeks after final vaccination, mice were infected by aerosol with approximately 100 colony forming units (CFU) of the virulent *M. tuberculosis* H37Rv strain (Fig. 1a). Bacterial load in the lungs was significantly reduced by both BCG and mRNA^CV2^ compared to PBS vaccination at 4 weeks post challenge (Fig. 2a). The protective effect induced by each of these vaccines was comparable, resulting in a 0.7 and 0.9 Log_10_ reduction in CFU, respectively. Protection afforded by mRNA^CV2^ did not extend to the spleen, suggesting limited efficacy against bacterial dissemination (Fig. 2b). In contrast, BCG vaccination significantly reduced the *M. tuberculosis* load in the spleen. The reduction of bacterial burden in the lungs was accompanied by a decrease in pathological changes observed in histological examination, further supporting pulmonary protection against *M. tuberculosis*. The lungs from PBS-vaccinated mice contained large areas of inflammatory infiltrate (Fig. 2c). BCG-vaccinated lungs had smaller lesions with less lung involvement, whilst the lowest level of cellular infiltration was observed in mRNA^CV2^-vaccinated mice. *M. tuberculosis*-challenged mice vaccinated with mRNA^CV2^ had a greater frequency of cytokine-secreting CD4^+^ T cells than mice vaccinated with either BCG or PBS (Fig S5a), notably the triple-positive polyfunctional population (Fig. 2d). IL-17A-producing CD4^+^ T cells were also elevated, although, as observed pre-vaccination, frequencies of this cell type were low (Fig. 2e). Notably, compared to BCG or PBS, mRNA-vaccination elicited Ag85B:I-A^b+^ CD4^+^ T cells that remained elevated post-infection (Fig. 2f). These findings were recapitulated in the dLN (Supplementary Fig. 5b-d). Hence, mRNA^CV2^ can significantly reduce virulent *M. tuberculosis* burden in the lungs of vaccinated mice, to a level at least as comparable to that of BCG, and this is associated with a strong vaccine-specific T cell response.

**Figure 2:**
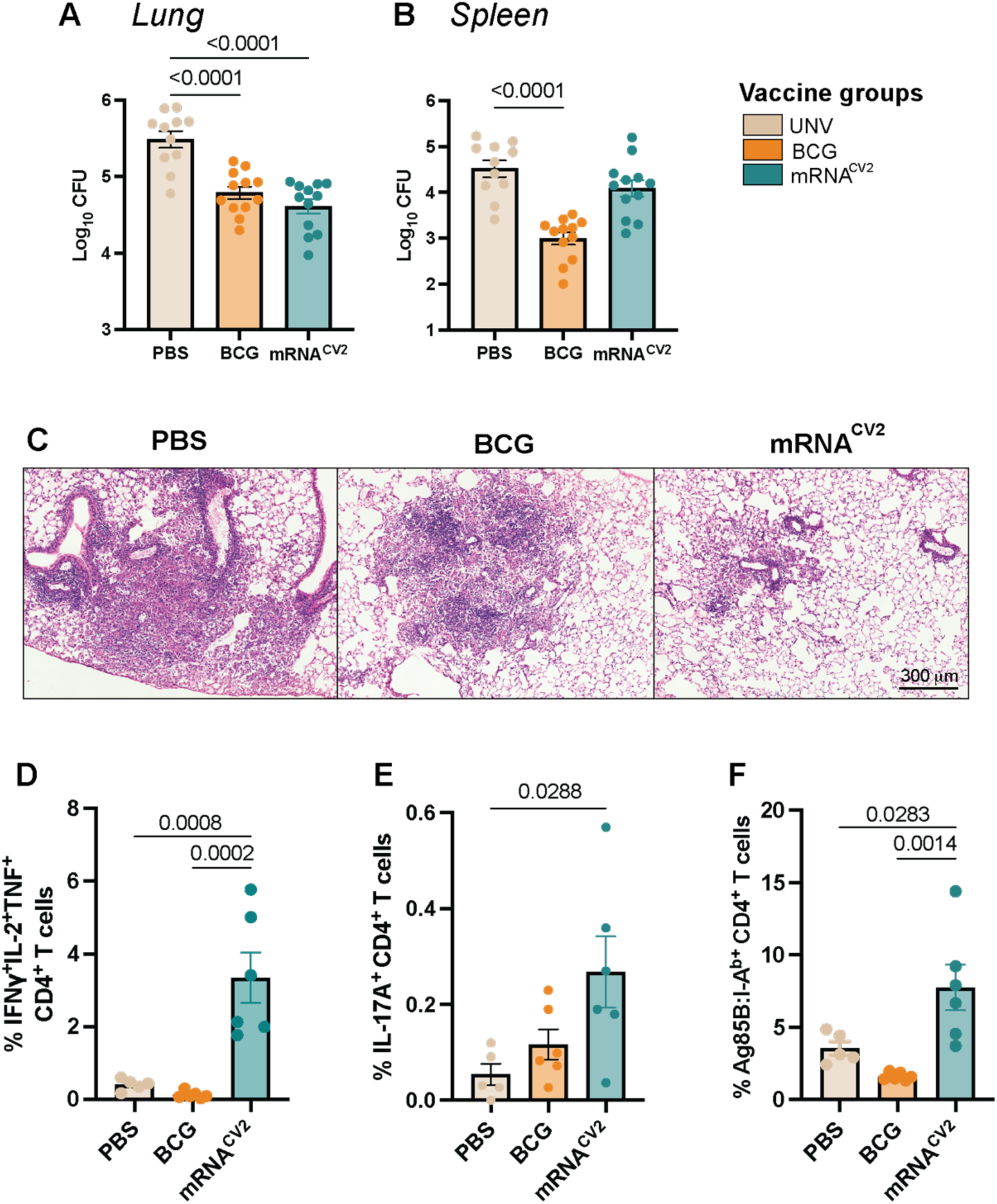
mRNA^CV2^ is protective in a mouse model of *M. tuberculosis* infection. *M. tuberculosis* CFU in **(A)** lung and **(B)** spleen of PBS, BCG or mRNA^CV2^-vaccinated mice 4 weeks after aerosol challenge with ∼100 CFU of *M. tuberculosis*. **(C)** Representative areas of haematoxylin and eosin (H&E) stained middle right lobes from PBS-, BCG-, and mRNA^CV2^-vaccined mice imaged at ×40 magnification. Proportions of CysVac2-specific **(D)** polyfunctional CD4^+^ T cells, **(E)** IL-17A-producing CD4^+^ T cells, and **(F)** Ag85B:I-A^b^ tetramer^+^ CD4^+^ T cells in the lung. Protection data **(A, B)** are pooled from 2 independent experiments and expressed as Log_10_ of mean of CFU ± SEM. Immunogenicity data **(D-F)** are expressed as mean +/-SEM with 5-6 mice in each group and representative of 2 independent experiments. Significance determined by **(A, B)** One-way ANOVA with Dunnett’s *post-hoc* test, and **(D-F)** One-way ANOVA corrected with Tukey’s *post-hoc* test.

### Intramuscular vaccination with mRNA^CV2^ induces rapid recruitment of immune cells to the draining lymph node and lung

We next profiled the spatiotemporal dynamics of immune cell recruitment by intramuscular mRNA^CV2^ vaccination to determine how these parameters correlated with the protective immune responses observed. C57BL/6 mice were vaccinated with a single dose of either PBS or mRNA^CV2^ and immune cell populations in the draining lymph node (dLN) and lung were assessed by flow cytometry (Supplementary Fig. 4) at timepoints outlined in Fig. 3a. In the dLN there was a rapid influx of neutrophils that peaked at 6 hours after vaccination with mRNA^CV2^ and declined after 1 day (Fig. 3b). Ly6C^+^ monocytes (Fig. 3c) and macrophages (Fig. 3d) peaked at 1-day post-vaccination; however, both subsets returned to baseline by day 7. Conventional dendritic cell populations 1 and 2 (cDC1 and cDC2, respectively) were both elevated in the dLN following mRNA^CV2^ vaccination (Fig. 3e-g). Migratory and resident subsets were defined by differential expression of CCR7, a chemokine receptor that directs migration of cells from peripheral tissues to the dLN^23,24^. Migratory DCs were the dominant phenotype of both cDC populations, appearing first in large numbers at day 1-2 and remaining elevated until day 5. Resident cDC1 and cDC2 populations were more stable, with a slight increase at day 2-3 that returned to baseline by day 7-10. Both B cells and CD4^+^ T cells peaked relatively early at day 2 or 1 respectively (Fig 4a, c), whilst CD8^+^ T infiltration into the dLN was delayed, peaking at day 5 post-immunisation (Fig. 4b). Vaccine-specific Ag85B:I-A^b+^ CD4^+^ T cells peaked only at 7 days post-vaccination before contracting to baseline by Day 14 (Fig. 4d, f). These Ag85B:I-A^b+^ CD4^+^ T cells exhibited a Th1-like phenotype, as determined by expression of CXCR3 (Fig. 4g). Additionally, relative to total CD4^+^ T cells, the tetramer-positive subset was enriched for expression of CD11a and PD-1 (Fig. 4e).

**Figure 3:**
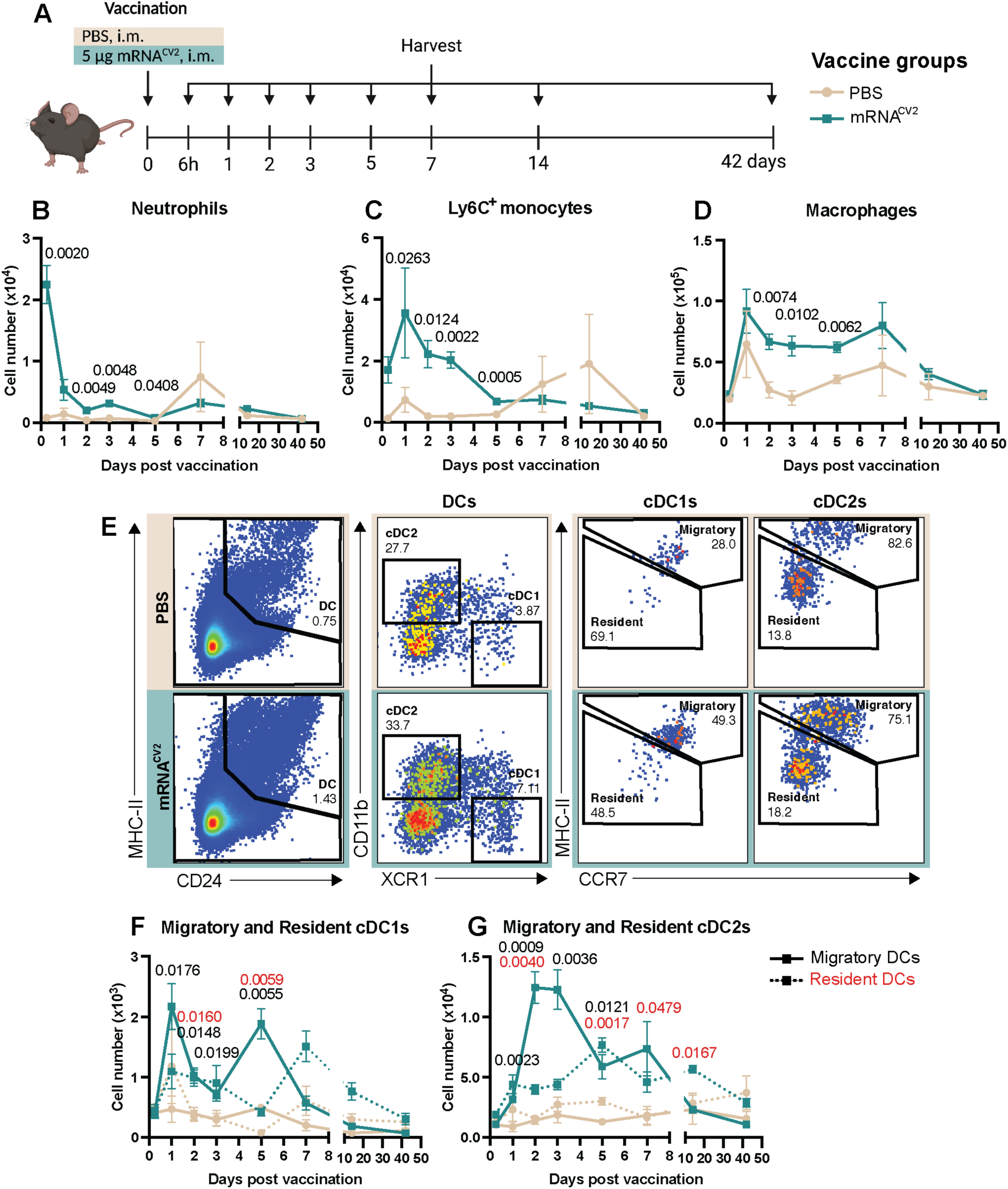
Innate immune cell dynamics in the inguinal lymph node after a single intramuscular dose of mRNA^CV2^. **(A)** Schematic of experimental outline. **(B-G)** Cell number of innate immune cell subsets at 6 hours, 1, 2, 3, 5, 7, 14 and 42 days after PBS or mRNA^CV2^ vaccination. **(E)** Representative cytometric plots of total dendritic cells (DCs) conventional DC subsets 1 and 2, and migratory and resident DCs at day 5. **(F, G)** For DC subsets, migratory and resident populations are denoted by solid and dashed lines respectively. Data are expressed as mean ± SEM with 3-4 mice in each vaccine group and representative of 2 independent experiments for all timepoints except 42 days. Significance between groups was determined by multiple unpaired t-tests. In **(E, F)** significance between vaccine groups for migratory and resident DCs is denoted in black and red, respectively.

**Figure 4:**
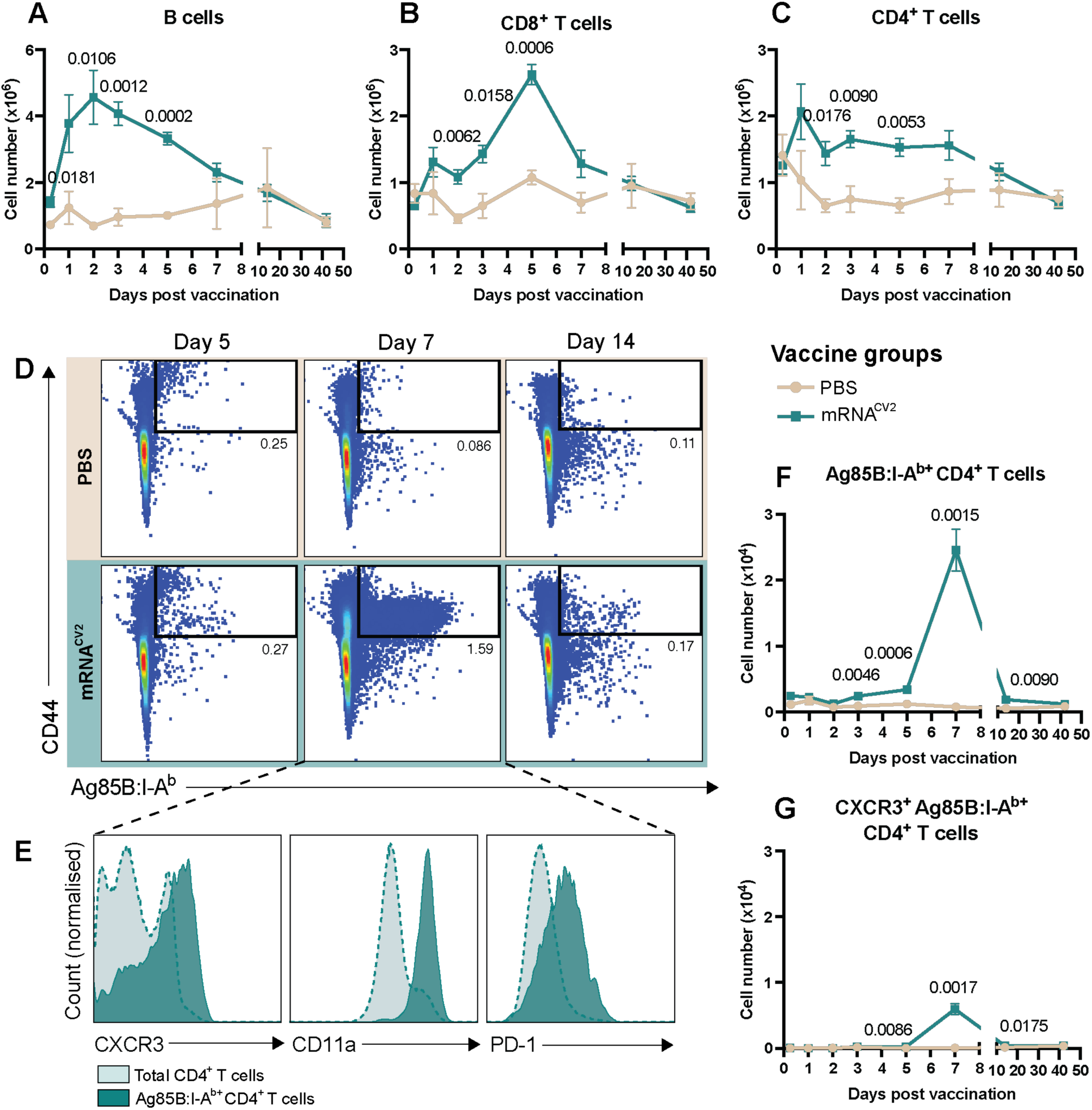
Adaptive immune cell dynamics in the inguinal lymph node after a single intramuscular dose of mRNA^CV2^. **(A-G)** Cell number of adaptive immune cell subsets at 6 hours, 1, 2, 3, 5, 7, 14 and 42 days after PBS or mRNA^CV2^ vaccination. **(D)** Representative cytometric plots of CD4^+^ T cells expressing the Ag85B:I-A^b^ tetramer at 5, 7 and 14 days after vaccination. **(E)** Representative histograms of CXCR3, CD11a and PD-1 expression by total or Ag85B:I-A^b+^ CD4^+^ T cells at day 7. Data are expressed as mean ± SEM with 3-4 mice in each vaccine group and representative of 2 independent experiments for all timepoints except 42 days. For **(E)**, counts were normalised to mode. Significance between groups was determined by multiple unpaired t-tests.

Given the lung is the primary site of *M. tuberculosis* infection, immune responses in this organ were also examined following mRNA^CV2^ vaccination. Early peaks of neutrophils, Ly6C^+^ monocytes and macrophages were observed after 6 hours, but cell number contracted rapidly to the level of PBS-vaccinated controls (Supplementary Fig. 6a-e). There was no increase in the B or T cell compartments (Supplementary Fig. 7a-c); however, Ag85B:I-A^b+^ CD4^+^ T cells were detected from 5 days after vaccination (Supplementary Fig. 7d, e). As in the dLN, these cells were of a Th1-type phenotype (Supplementary Fig. 7g) and expressed PD-1 and CD11a (Supplementary Fig. 7e), the latter of which is a key receptor involved in the homing of T cells to the lung ^25^. Overall, these data indicate that a single intramuscular dose of mRNA^CV2^ is immunogenic and capable of inducing a rapid vaccine-specific response both locally and systemically in mice.

### mRNA^CV2^ enhances immune responses and confers long-term protection against M. tuberculosis in BCG-primed mice

The high global coverage of infant BCG immunisation^26^ necessitates an assessment of the ability of TB vaccine candidates to boost pre-existing immunity afforded by BCG. To assess this, C57BL/6 mice were primed with BCG 6 months prior to boosting with 2 doses of the mRNA^CV2^ vaccine, 3-weeks apart (Fig. 5a). This boosting regimen was compared to age-matched mice who had received BCG alone, mRNA^CV2^ alone, or no vaccination. Prior to boosting, there were negligible residual CD4^+^ T cell responses in BCG-vaccinated mice above unvaccinated mice (Fig. 5b). A week after the second booster dose, there was a significant increase in vaccine-specific immune responses for both groups who received mRNA^CV2^ (Fig. 5b). At this post-boost timepoint there were significantly greater frequencies of triple-positive (IFNγ^+^, IL-2^+^, TNF^+^) CD4^+^ T cell populations in mice who received mRNA^CV2^ compared to those who only received BCG (Fig. 5b). This difference was most apparent when examining the frequency of IFNγ^+^ and TNF^+^ CD4^+^ T cells, which were significantly elevated in the BCG-mRNA^CV2^ group compared to those who received BCG or mRNA^CV2^ alone (Fig. 5c).

**Figure 5:**
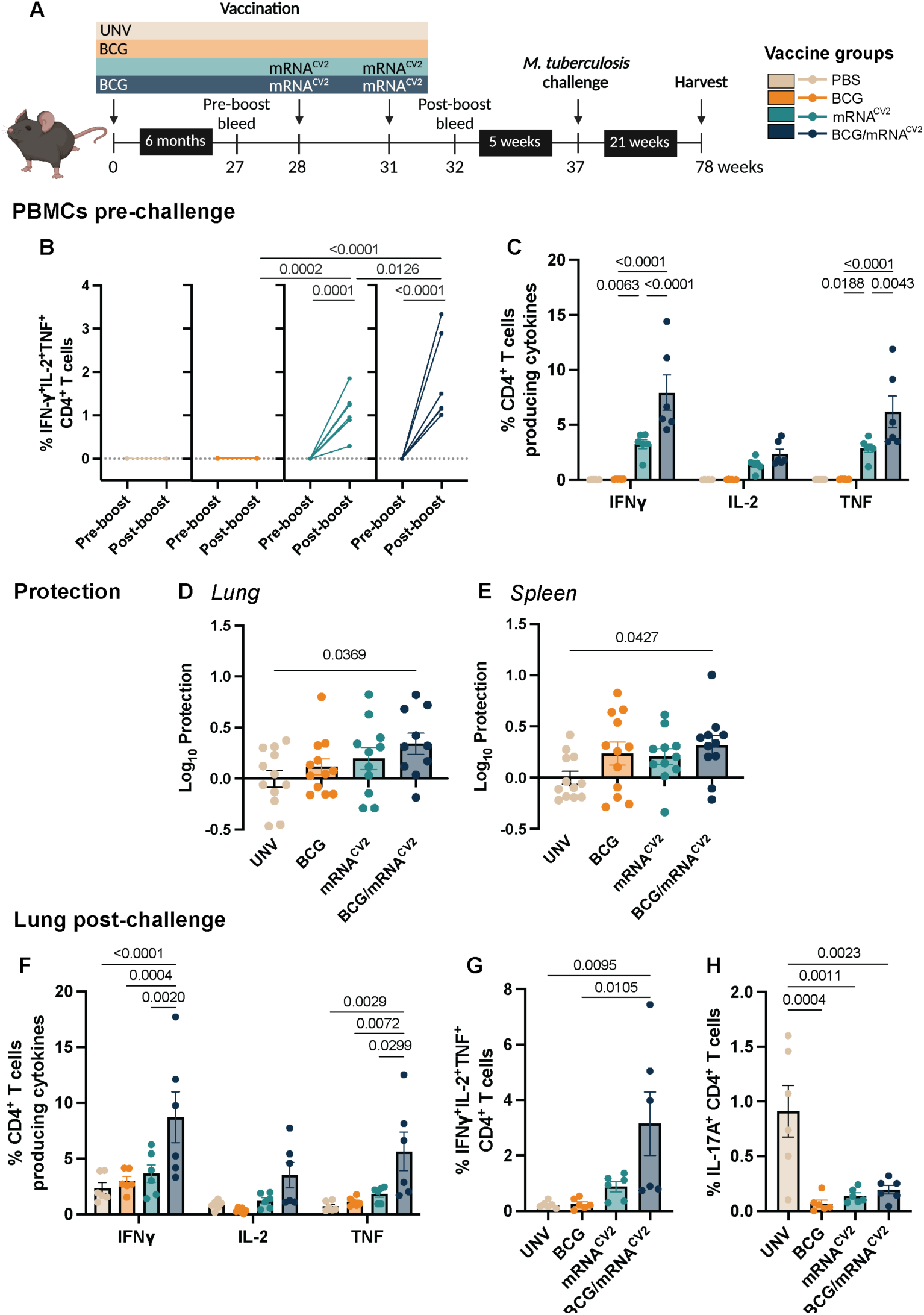
mRNA^CV2^ is effective as a booster vaccination for BCG against *M. tuberculosis*. **(A)** Schematic of experimental outline. **(B)** Proportions of CysVac2-specific polyfunctional CD4^+^ T cells in PBMCs of immunised C57BL/6 mice at 27 (pre-boost) or 32 weeks (post mRNA boost). **(C)** Proportions of IFNγ-, IL-2- and TNF-producing CD4^+^ T cells in peripheral blood 32 weeks after initial vaccination. Log_10_ of delta protection from *M. tuberculosis* in **(D)** lung and **(E)** spleen. Proportions of **(F)** single-cytokine secreting, **(G)** polyfunctional, or **(H)** IL-17A-producing CD4^+^ T cells in the lungs of vaccinated mice 21 weeks after aerosol challenge with ∼100 CFU of *M. tuberculosis*. Immunogenicity data **(B, C, F-H)** are expressed as mean ± SEM with 5-6 mice in each group and representative of 2 independent experiments. Protection data **(D, E)** are pooled from 2 independent experiments and expressed as the mean of Log_10_ CFU ± difference of each individual mouse compared to the mean of unvaccinated mice. Significance determined by **(B)** Two-way ANOVA, **(C, F)** Two-way ANOVA with Tukey’s *post-hoc* test, **(D, E)** One-way ANOVA with Dunnett’s *post-hoc* test, and **(G, H)** One-way ANOVA with Tukey’s *post-hoc* test.

To determine if mRNA^CV2^ boosting of BCG could improve protection against *M. tuberculosis* infection, 6 weeks after the final vaccination mice were challenged and bacterial load examined after 21 weeks, at which timepoint the effect of BCG protection wanes in mice^17,27^. To account for variation between experiments, data were normalised to represent the Log_10_ difference in bacterial load of vaccinated mice compared with the mean of unvaccinated mice in each experiment. Relative to unvaccinated mice, mice immunised with both BCG and mRNA^CV2^ were the only group to exhibit significantly increased protection in either the lung (Fig. 5d) or spleen (Fig. 5e). Neither BCG nor mRNA^CV2^ alone groups conferred statistically significant protection against infection. CysVac2 restimulation of lung single-cell suspensions revealed that only BCG-primed mRNA^CV2^-boosted mice had significantly greater IFNγ- and TNF-secreting, as well as triple-positive (IFNγ^+^, IL-2^+^, TNF^+^) CD4^+^ T cell populations (Fig. 5f, g), compared to other groups. This was also observed in the dLN (Supplementary Fig. 8). Protection was associated with a decreased frequency of IL-17A^+^ CD4^+^ T cells in the lung for all vaccinated groups (Fig. 5h). Overall, these data indicate that mRNA^CV2^ delivery is suitable for boosting protection afforded by the standard of care BCG vaccine, with both vaccine-specific immunity and protection enhanced when combined with BCG.

## Discussion

The continued absence of a candidate to replace the only approved TB vaccine, BCG, necessitates further research into alternative vaccine approaches^1^. Here we describe the first report of a highly protective LNP-mRNA vaccine for pulmonary TB in mice, expressing a fusion of two *M. tuberculosis* antigens. The protection observed here exceeds that seen in the small number of previous reports using RNA-based vaccines for TB^28–30^ (Fig. 2). This may be due to our use of a highly immunoprotective antigen combination ^17–20^, together with an immunogenic LNP formulation^21^. Protection in our study was associated with a high frequency of vaccine-specific CD4^+^ T cells producing IFNγ, IL-2 and TNF in mice immunised with mRNA^CV2^; both before and after aerosol infection with *M. tuberculosis* (Fig. 1, 2). These polyfunctional CD4^+^ T cells are frequently correlated with protection against *M. tuberculosis* in other mouse and non-human primate studies ^17^, although this has not always recapitulated in humans^31^. The frequency of these cytokine-producing CD4^+^ T cells, as well as antigen-specific Ag85B:I-A^b+^ CD4^+^ T cells, was far greater than we have previously observed following vaccination with an adjuvanted form of the CysVac2 protein^17,18^. This is presumably due to the mRNA vaccine platform, which is known to produce strong Th1-type CD4^+^ T cell responses in mice^32,33^, potentially through the induction of Type I IFNs^33–36^. Whether this strong T cell immunity mediated by TB mRNA vaccines will translate to humans is yet to be determined.

In addition to Th1 CD4^+^ T cell responses following mRNA^CV2^ vaccination, there was an increase in IL-17A-producing CD4^+^ T cells in the circulation (Fig. 1) and lungs of vaccinated mice during acute infection (Fig. 2), albeit at low frequencies. We have previously observed vaccine-mediated induction of IL-17A by Th17 CD4^+^ T cells to correlate with greater protection against *M. tuberculosis* in mice at early stages of infection^17,19,20^. Intriguingly, here IL-17A was associated with decreased protection during chronic *M. tuberculosis* infection (Fig. 5), suggesting excessive induction of this cytokine long-term may be detrimental at later stages of infection^37^. Regardless, pulmonary administration of BCG has been shown to enhance the development of protective IL-17A^+^ CD4^+^ T cells compared to subcutaneous immunisation^38,39^. This is also the case for intratracheal, rather than intramuscular, administration of CysVac2 subunit vaccines^18,20^. Current standard LNP formulations induce highly inflammatory responses that are damaging to the lung if delivered directly, making them unsuitable for mucosal delivery^40^. Development of LNP-mRNA formulations that can be safely delivered to the lung, either directly or with pulmonary targeting lipids^41^, to enhance Th17 CD4^+^ T cell responses early post-vaccination is of interest. This may serve to increase localised immune responses and improve protection against respiratory pathogens.

Intramuscular vaccination with a single dose of mRNA^CV2^ induced a rapid, transient influx of neutrophils, followed by Ly6C^+^ monocytes, macrophages and migratory DCs, to the dLN (Fig. 3). This acute, robust inflammatory response has previously been reported to be mediated by the ionisable lipid component of standard LNP formulations, and may explain the effectiveness of the mRNA vaccine platform^40^. Translation of antigen from the mRNA construct is predominantly performed by these APCs, which subsequently present said antigen to naïve CD4^+^ T cells^35,42^. Hence, their rapid infiltration to the lymph node observed here following mRNA^CV2^ vaccination is beneficial to a rapid immune response. Whilst migratory DCs travel from peripheral tissues to the dLN following activation, resident DC populations generally remain stable^43,44^. In addition to an early influx of migratory DCs in mRNA^CV2^-vaccinated mice, we also observed a slight delayed increase in the resident DC population. This could be result of a loss of CCR7 expression from migratory DCs after arriving in the lymph node^45^. Regardless, both subsets can individually present antigen to T cells, and migratory DCs may also transfer captured antigen to lymph node-resident DCs to enhance T cell activation^46,47^. Of the other DC subsets we investigated, cDC1s preferentially activate CD8^+^ T cells^48^, whilst cDC2s primarily present antigens to CD4^+^ T cells^49^. Recruitment of both these subsets to the dLN suggests the response following vaccination is well balanced. Indeed, we observed similar dynamics of CD4^+^ and CD8^+^ T cells recruited to the dLN (Fig. 4). One limitation of this study is the lack of a defined CD8^+^ T cell epitope on the CysVac2 antigen in C57BL/6 mice, which limits analysis of the recruitment and phenotype of antigen-specific CD8^+^ T cells induced by the vaccine. Nonetheless, mRNA^CV2^ immunisation established an environment in the dLN for the recruitment and activation of adaptive immunity, notably, tetramer-positive CD4^+^ T cells exhibiting a Th1-like phenotype (Fig. 4).

Unlike in the dLN, we did not observe sustained infiltration of innate immune cells into the lung (Supplementary Fig. 6), possibly reflecting the lack of initial priming of adaptive immunity in the lung following intramuscular routes of vaccination^50^. Despite this, we observed the establishment of a Th1-type Ag85B:I-A^b+^ CD4^+^ T cell population (Supplementary Fig. 7). These cells expressed phenotypic markers of lung tissue homing and accessory proteins associated with resident memory T cells (T_RM_)^51^, which facilitate rapid clearance of bacteria and enhance protection against *M. tuberculosis*^39,52,53^. At this early timepoint, however, they may not represent true T_RM_ cells. Most studies of currently approved mRNA formulations for COVID-19 demonstrate an inability to establish CD4^+^ T_RM_ cell populations in both mice and humans^33,54^. CD8^+^ T_RM_ cells have been observed in mice using mRNA encoding the influenza nucleoprotein encapsulated in nanoparticles^55^, but not using LNP-formulated SARS-CoV-2 spike antigen^56^, suggesting this outcome could be antigen and/or formulation dependent. Notably, despite strong protection in the lung, the ability of mRNA^CV2^ to protect against disseminated infection was limited. Protection in the spleen has previously been observed with protein subunit formulations of CysVac2^17–20^, suggesting this is not the result of antigen selection, but likely the LNP-mRNA platform. Thus, it is possible that intramuscular mRNA vaccination may not induce sufficient lung resident T cells to prevent early dissemination of *M. tuberculosis* bacilli following infection. Mucosal delivery or organ targeting may present a strategy to promote the LNP-mRNA-mediated development of CD4^+^ T_RM_ cells in the lungs^20,55^.

Novel TB vaccines must be effective as a booster vaccination to BCG as the majority of the global population has received the vaccine as part of childhood immunisation programs because of its documented protective effect against TB meningitis and disseminated TB in children^26^. BCG also induces non-specific immunity and reduces mortality against unrelated respiratory pathogens^57^. In this study, we demonstrated that boosting BCG-primed mice with mRNA^CV2^ enhanced the antigen-specific immune response above that observed with mRNA^CV2^ or BCG alone (Fig. 5). This was associated with protection against chronic *M. tuberculosis* infection in the lung and protection from dissemination to the spleen, which was not observed in mice vaccinated with either component individually. Following vaccination, BCG is rapidly controlled by the host immune response during a period of active replication, so is unable to induce immunity against antigens expressed during latent stages of *M. tuberculosis* infection^58^. We propose that inclusion of latency antigen, CysD, in mRNA^CV2^ may be responsible for enhancing protection in BCG-primed mice at this extended timepoint of chronic infection. Furthermore, the inability to generate enduring protective immunity is an established challenge for mRNA vaccines^59^. Heterologous prime-boost immunisation regimens have proven superior in COVID-19 with the ChAdOx1 recombinant viral and BNT162b2 mRNA vaccines^60^. The observed synergistic effect of combining the two distinct vaccine platforms of BCG and mRNA here supports a strategy by which to improve the efficacy and durability of mRNA vaccines.

In summary, we have demonstrated that the LNP-mRNA platform is both immunogenic and protective in a murine model of *M. tuberculosis* infection. This approach holds significant potential in the fight against TB, both as a standalone vaccine, and as a booster to the existing BCG vaccine. The promising results of this study underscore the need for further research and clinical trials to complement existing efforts^6^ to explore the full capability of mRNA vaccines for combating TB.

## Data Availability

The data that support the findings of this study are available from the corresponding author upon reasonable request.

### Acknowledgments

This work was supported by the NHMRC Centre of Research Excellence in Tuberculosis Control (APP1153493), and MRFF mRNA Clinical Trial Enabling Infrastructure grant (MRFCTI000006). We thank the NIH Tetramer Core Facility (Atlanta, USA) for providing the Ag85B_240-254_:I-A^b^ Class II MHC tetramer.

## Author contributions

HL: Conceptualisation, Experimental work, Data curation, Formal analysis, Methodology, Writing— original draft. HAW: Conceptualisation, Experimental work, Methodology, Writing—Review and Editing. SAB: Conceptualisation, Experimental work, Methodology, Writing—Review and Editing. LL: Experimental work, Methodology, Data curation. TW: Experimental work, Methodology, Data curation, Formal analysis, Writing—Review and Editing. WJB: Conceptualisation, Writing – Review and Editing. MS: Conceptualisation, Methodology, Writing—Review and Editing. CWP: Conceptualisation, Methodology, Writing—Review and Editing, Funding Acquisition. JAT: Conceptualisation, Formal analysis, Methodology, Writing—Review and Editing, Funding Acquisition. CC: Conceptualisation, Methodology, Writing—Review and Editing, Funding Acquisition. All authors have read and agreed to the published version of the manuscript.

