## Supporting information for "A lipid nanoparticle-mRNA vaccine provides potent immunogenicity and protection against *Mycobacterium tuberculosis*"

**Lukeman *et al.***

**Supplementary Table 1:** Monoclonal antibodies used for flow cytometry

| Antibody | Clone | Dilution | Manufacturer |
|---|---|---|---|
| Anti-mouse B220 AF594 | RA3-6B2 | 1:200 | BioLegend |
| Anti-mouse B220 BUV395 | RA3-6B2 | 1:200 | BD Horizon |
| Anti-mouse B220 BUV737 | RA3-6B2 | 1:200 | BD Horizon |
| Anti-mouse CCR7 BV605 | 4B12 | 1:200 | BioLegend |
| Anti-mouse CD103 BV786 | M290 | 1:200 | BioLegend |
| Anti-mouse CD11a BV510 | M17/4 | 1:200 | BD Biosciences |
| Anti-mouse CD11b APC-Cy7 | M1/70 | 1:200 | BD Pharmingen |
| Anti-mouse CD11c AF647 | N418 | 1:200 | BD Pharmingen |
| Anti-mouse CD19 BV786 | 1D3 | 1:200 | BD Horizon |
| Anti-mouse CD24 BUV737 | M1/69 | 1:200 | BD Biosciences |
| Anti-mouse CD25 FITC | 3C7 | 1:200 | BioLegend |
| Anti-mouse CD3 PE-Cy7 | 145-2C11 | 1:200 | BD Pharmingen |
| Anti-mouse CD4 AF700 | RM4-5 | 1:200 | BD Pharmingen |
| Anti-mouse CD44 BV605 | IM7 | 1:200 | BD Horizon |
| Anti-mouse CD44 FITC | IM7 | 1:200 | BD Pharmingen |
| Anti-mouse CD45 BV510 | 30-F11 | 1:200 | BD Horizon |
| Anti-mouse CD62L APC-Cy7 | MEL-14 | 1:200 | BioLegend |
| Anti-mouse CD62L BV650 | MEL-14 | 1:200 | BD Horizon |
| Anti-mouse CD64 PE-Cy7 | X54-5/7.1 | 1:200 | BioLegend |
| Anti-mouse CD69 FITC | H1.2F3 | 1:200 | BD Pharmingen |
| Anti-mouse CD8 APC-Cy7 | 53-6.7 | 1:200 | BD Pharmingen |
| Anti-mouse CD8 FITC | 53-6.7 | 1:200 | BD Pharmingen |
| Anti-mouse CD8 Pacific Blue | 53-6.7 | 1:200 | BD Pharmingen |
| Anti-mouse CXCR3 PerCP-Cy5.5 | CXCR3.173 | 1:200 | BioLegend |
| Anti-mouse F4/80 PE | BM8 | 1:200 | BioLegend |
| Anti-mouse FOXP3 PE-Cy7 | FJK-16s | 1:100 | Invitrogen |
| Anti-mouse GATA3 BUV395 | L50-823 | 1:100 | BD Horizon |
| Anti-mouse IFNγ PE-Cy7 | XMG1.2 | 1:200 | BD Pharmingen |
| Anti-mouse IL-17A BV421 | TC11-18H10 | 1:200 | BD Horizon |
| Anti-mouse IL-2 PE | JES6-5H4 | 1:200 | BD Pharmingen |
| Anti-mouse IL-4 PE-CF594 | 11B11 | 1:200 | BD Horizon |
| Anti-mouse IL-5 APC | TRFK5 | 1:200 | BD Pharmingen |
| Anti-mouse iNOS AF488 | CXNFT | 1:200 | Invitrogen |
| Anti-mouse Ki-67 PerCP-eF710 | SoLA15 | 1:400 | Invitrogen |
| Anti-mouse KLRG1 BV421 | 2F1 | 1:200 | BD Horizon |
| Anti-mouse Ly6C PerCP-Cy5.5 | HK1.4 | 1:200 | Invitrogen |

| | | | |
|---|---|---|---|
| Anti-mouse Ly6G BUV395 | 1a8 | 1:200 | BD Horizon |
| Anti-mouse MHC-II AF700 | M5-114.15.2 | 1:200 | BioLegend |
| Anti-mouse PD-1 BV711 | 29F.1A12 | 1:200 | BD Horizon |
| Anti-mouse PD-1 BV786 | 29F.1A12 | 1:200 | BD Horizon |
| Anti-mouse RORγT PE-CF594 | Q31-378 | 1:100 | BD Horizon |
| Anti-mouse Siglec F BV421 | E50-2440 | 1:200 | BD Pharmingen |
| Anti-mouse T-bet APC | 4B10 | 1:200 | BioLegend |
| Anti-mouse T-bet PE | 4B10 | 1:200 | Invitrogen |
| Anti-mouse TNF PerCP-Cy5.5 | MP6XT22 | 1:200 | BD Pharmingen |
| Anti-mouse XCR1 BV650 | ZET | 1:200 | BioLegend |

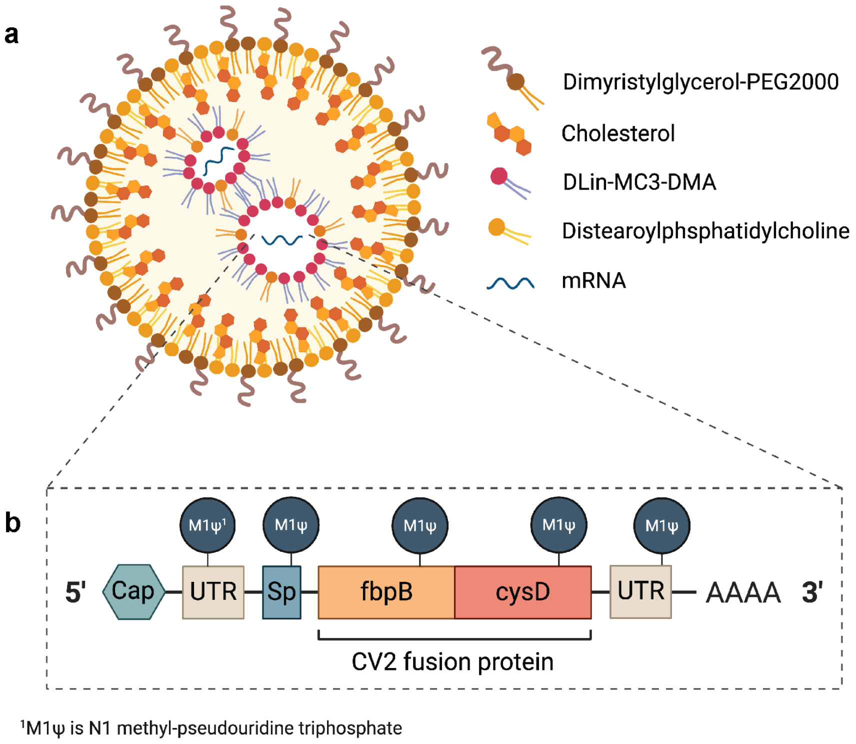


**Supplementary Figure 1: Design of mRNA$^{CV2}$ vaccine. (A)** Structure of the lipid nanoparticle delivery system. **(B)** mRNA construct contains TriLink CleanCap and 125 nucleotide polyA tail. Construct encodes 5' and 3'untranslated regions (UTR), SEAP signal peptide (Sp) and CysVac2 (CV2) fusion protein Antigen 85B (Ag85B) and Cystein. N1-methyl-pseudouridine triphosphate (M1ψ) is used instead of uridine triphosphate.

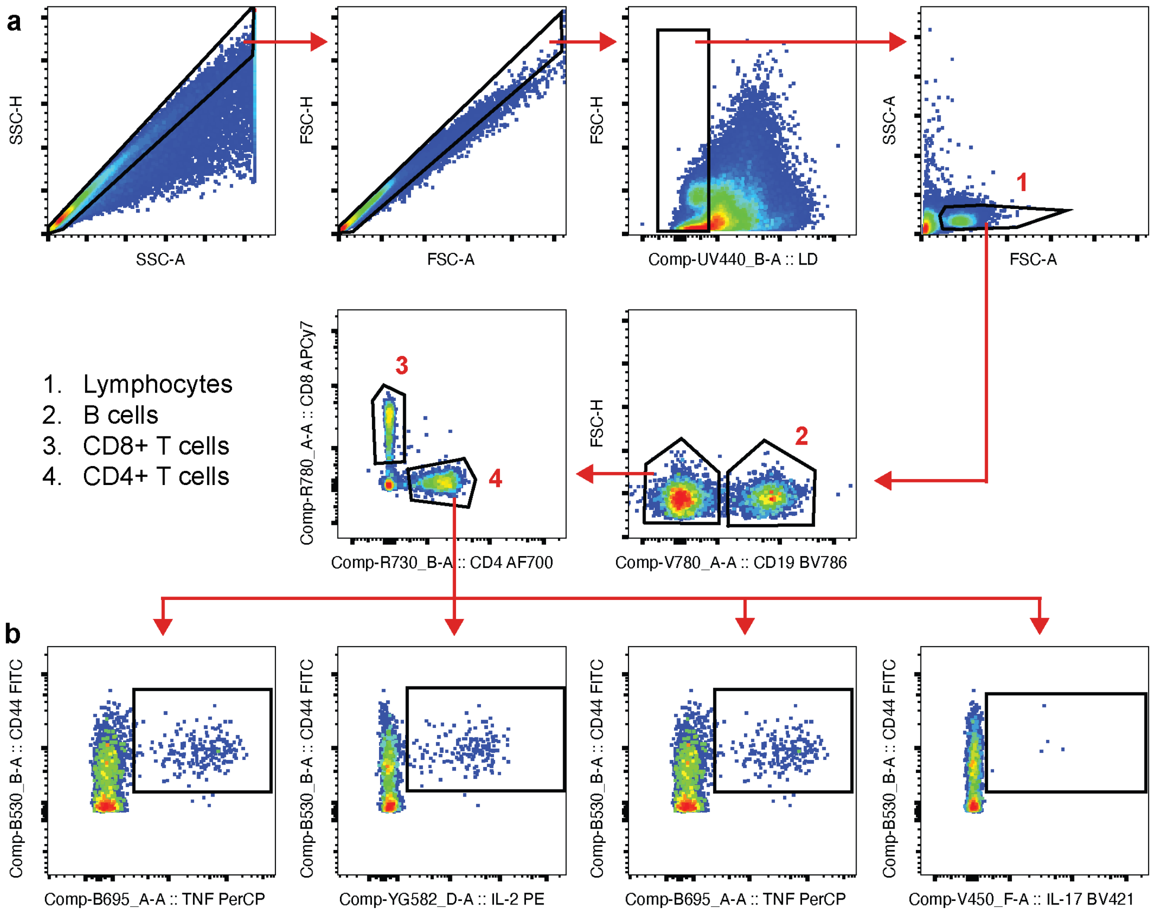


**Supplementary Figure 2: Manual gating strategy used for CysVac2 restimulation staining. (A)** Cells were gated by initially excluding doublets, then dead cells. Lymphocytes were selected by gating based on size. B cells were excluded by gating on CD19, then T cell subsets identified by expression of CD4 or CD8. **(B)** CD4$^{+}$ cytokine-expressing populations were identified.

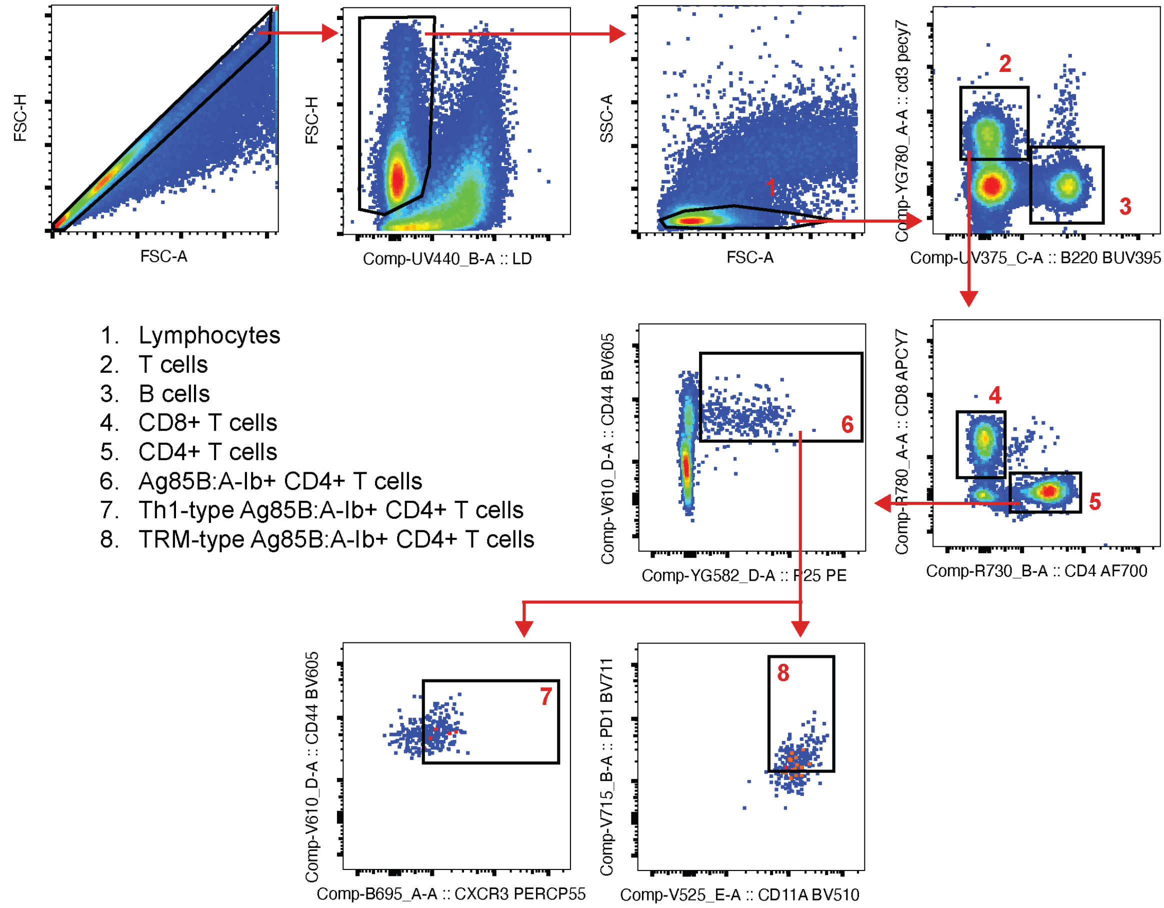


**Supplementary Figure 3: Manual gating strategy used for Tetramer staining.** Cells were gated by initially excluding doublets, then dead cells. Lymphocytes were selected by gating based on size. T cells and B cells were identified based on expression of B220 or CD3, then T cells were gated of CD4 or CD8 expression. Tetramer-specific CD4$^{+}$ T cells were identified by gating for Ag85B:A-I$^{b}$, then tissue resident memory- or Th1- type phenotypes were identified by gating for CXCR4 or CD11a and PD-1, respectively.

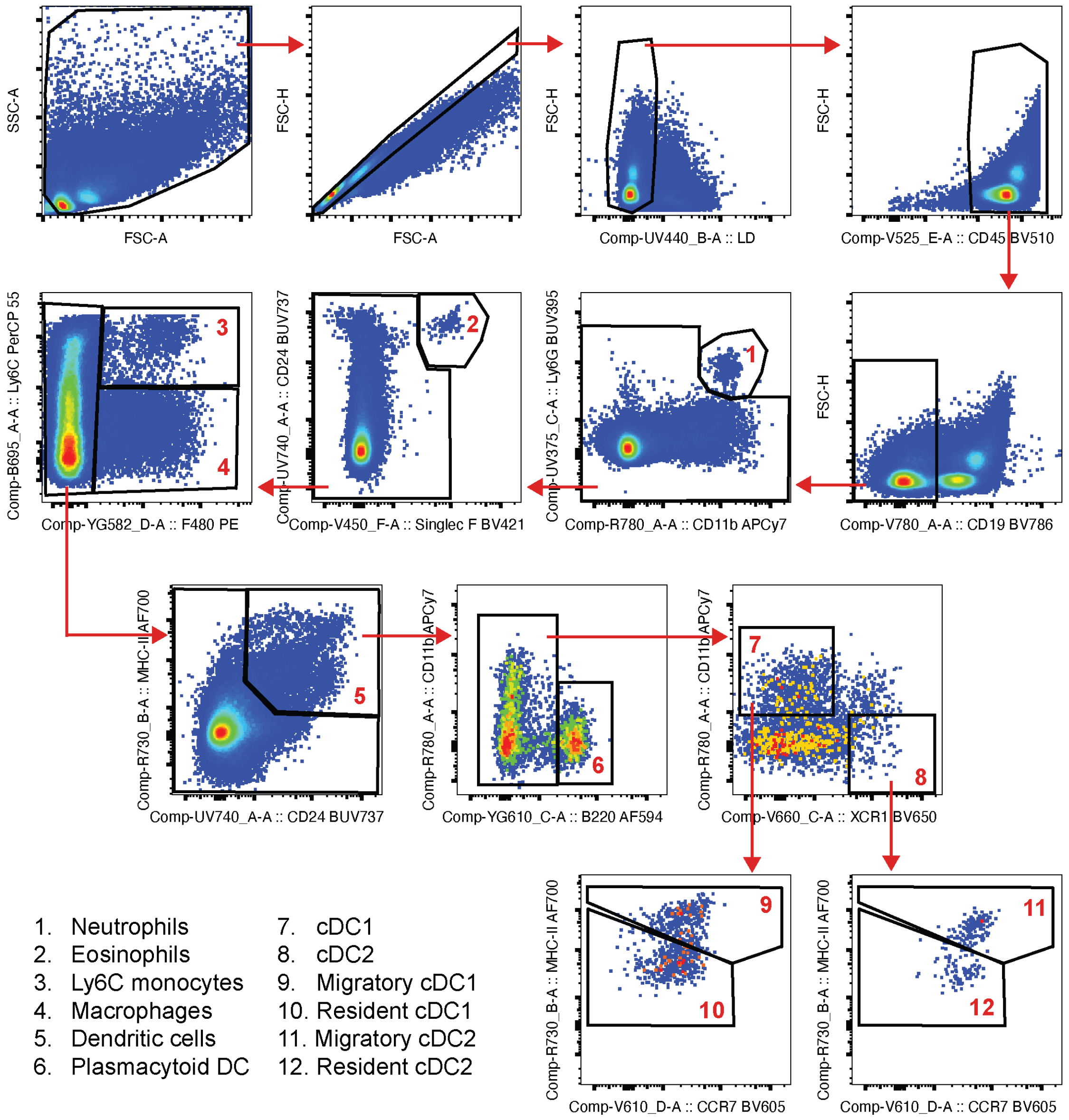


**Supplementary Figure 4: Manual gating strategy used for myeloid cells.** Cells were initially gated by excluding debris, doublets were excluded, then finally gated on live cells. Immune cells selected by gating on CD4, then myeloid cell subsets were individually identified in a stepwise manner.

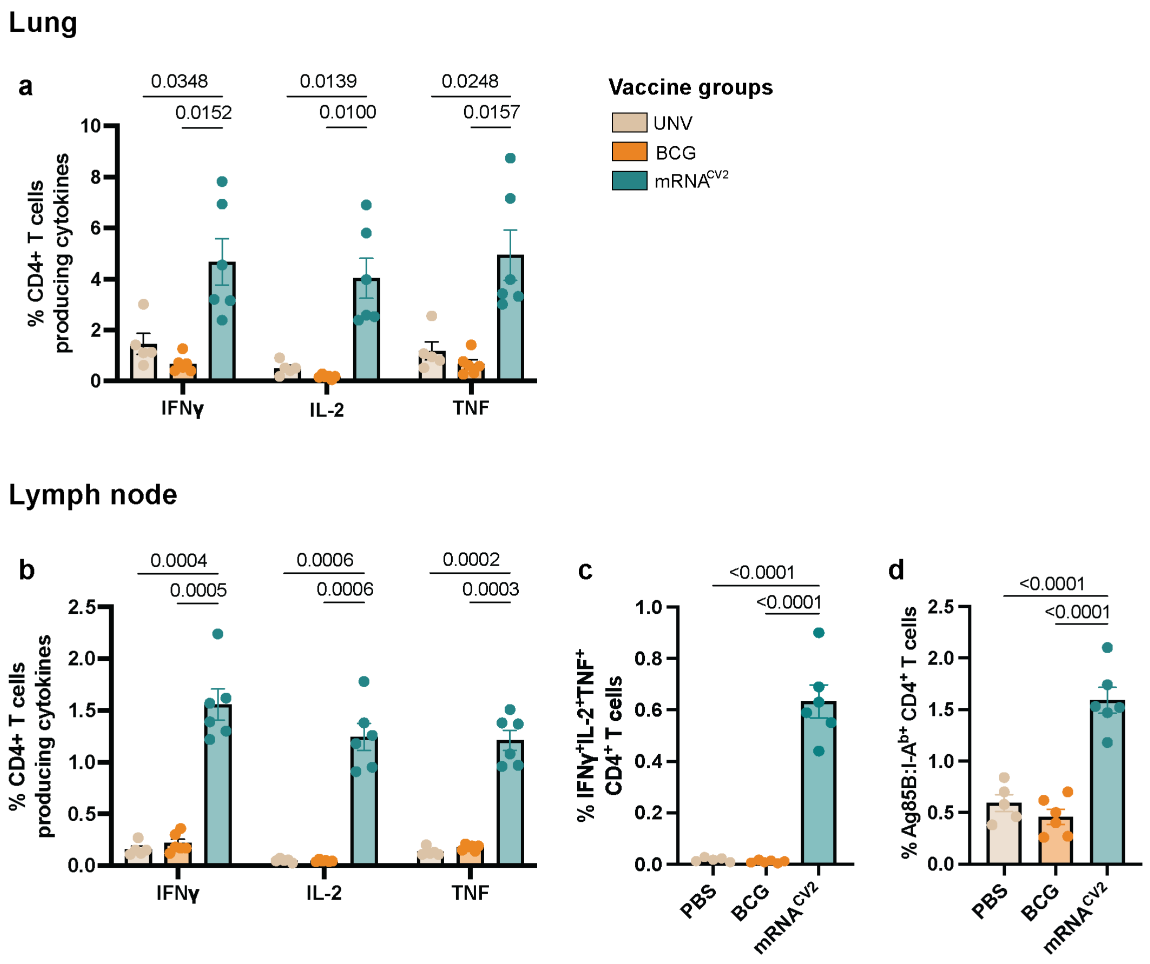


**Supplementary Figure 5: Potent CysVac2-specific CD4⁺ T cell responses following mRNA$^{CV2}$ vaccination in the lung and draining lymph node of *M. tuberculosis*-infected mice.** Proportions of CysVac2-specific IFNg-, TNF-, and IL-2-producing CD4$^+$ T cells in the **(A)** lung and **(B)** draining lymph node after M. tuberculosis infection. **(C)** Polyfunctional CD4$^+$ T cells and **(C)** Ag85B:I-A$^b$ tetramer$^+$ CD4$^+$ T cells in the lymph node. Data are expressed as mean +/- SEM with 5-6 mice in each group and representative of 2 independent experiments. Significance determined by **(A, C, D)** 1-way ANOVA with Tukey's post-hoc test and **(B)** 2-way ANOVA with Tukey's post-hoc test.

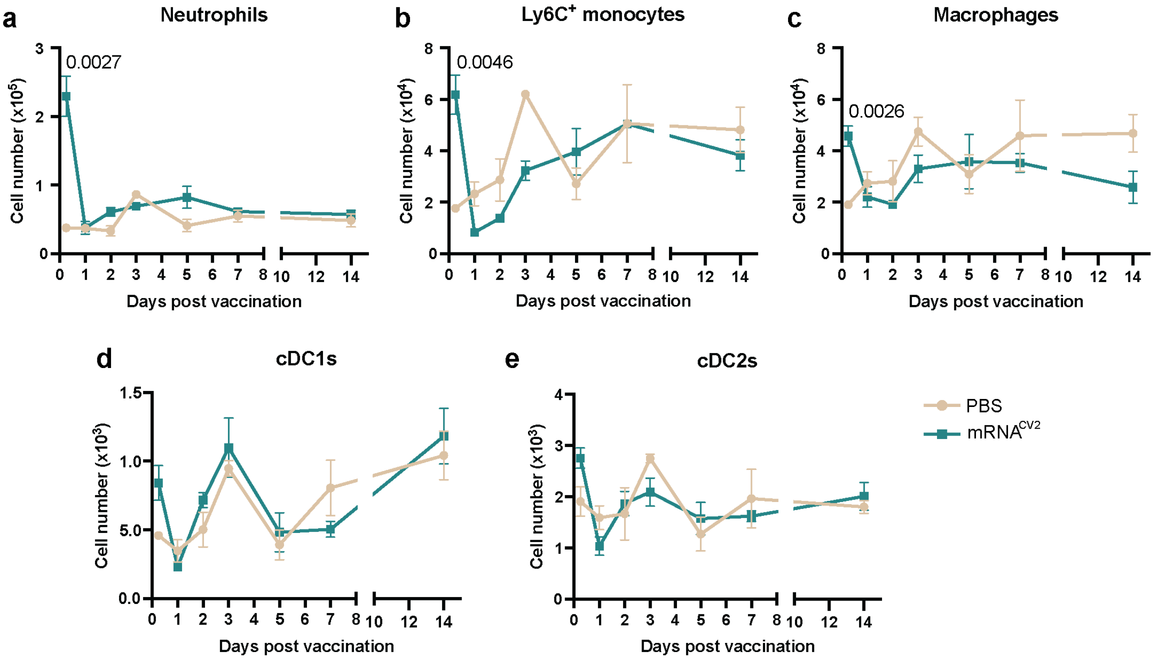


**Supplementary Figure 6: Innate immune cell dynamics in the lung after a single intramuscular dose of mRNA$^{CV2}$. (A-E)** Cell number of innate immune cell subsets at 6 hours, 1, 2, 3, 5, 7, 14 and 42 days after PBS or mRNA$^{CV2}$ vaccination. Data are expressed as mean ± SEM with 3-4 mice in each vaccine group and representative of 1 independent experiment. Significance between groups was determined by multiple unpaired t-tests.

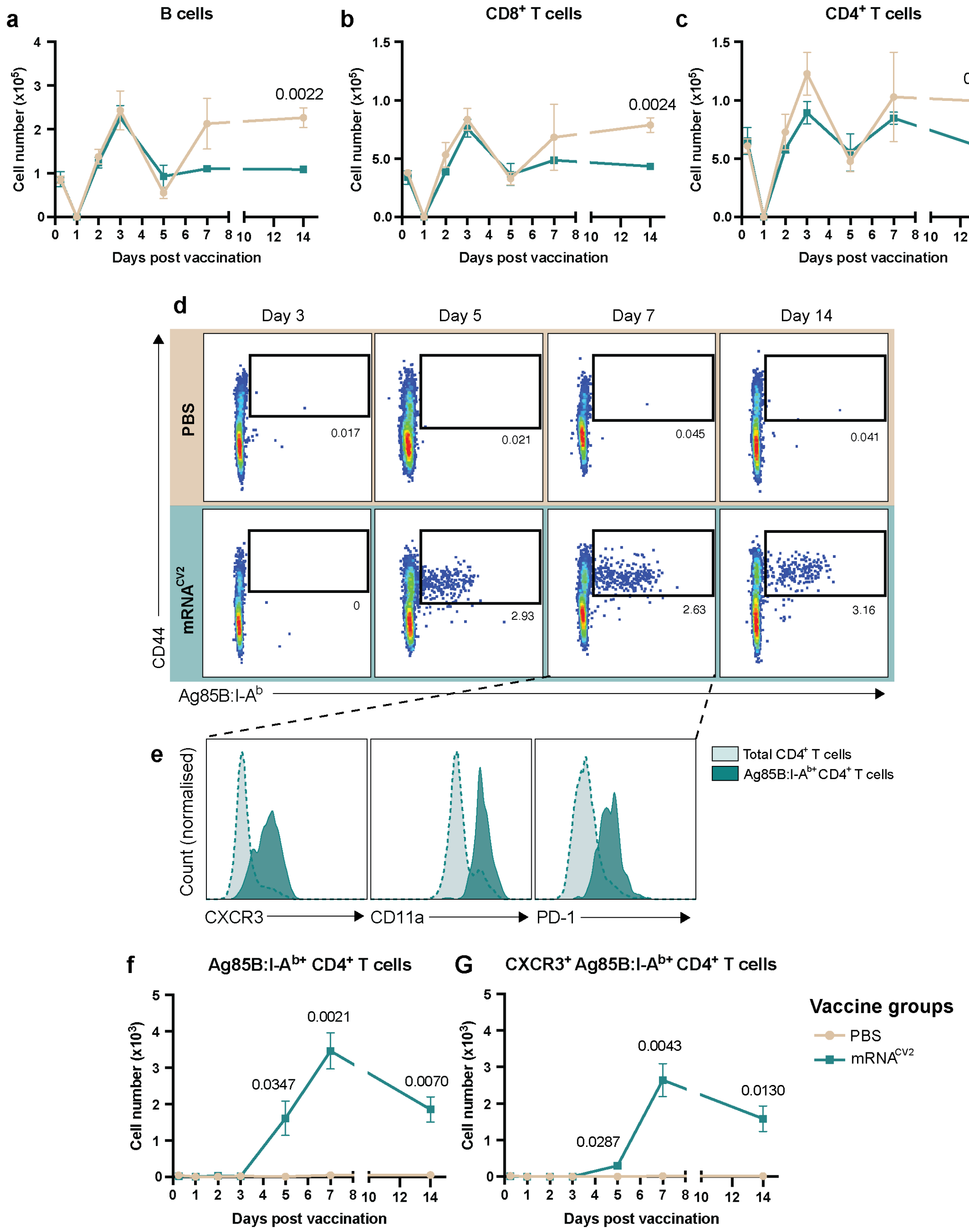


**Supplementary Figure 7: Adaptive immune cell dynamics in the inguinal lymph node after a single intramuscular dose of mRNA$^{CV2}$. (A-G)** Cell number of adaptive immune cell subsets at 6 hours, 1, 2, 3, 5, 7, 14 and 42 days after PBS or mRNA$^{CV2}$ vaccination. **(D)** Representative cytometric plots of CD4$^+$ T cells expressing the Ag85B:I-A$^b$ tetramer at 5, 7 and 14 days after vaccination. **(E)** Representative histograms of CXCR3, CD11a and PD-1 expression by total or Ag85B:I-A$^{b+}$ CD4$^+$ T cells at day 7. Data are expressed as mean ± SEM with 3-4 mice in each vaccine group and representative of 1 independent experiment. For **(E)**, counts were normalised to mode. Significance between groups was determined by multiple unpaired t-tests.

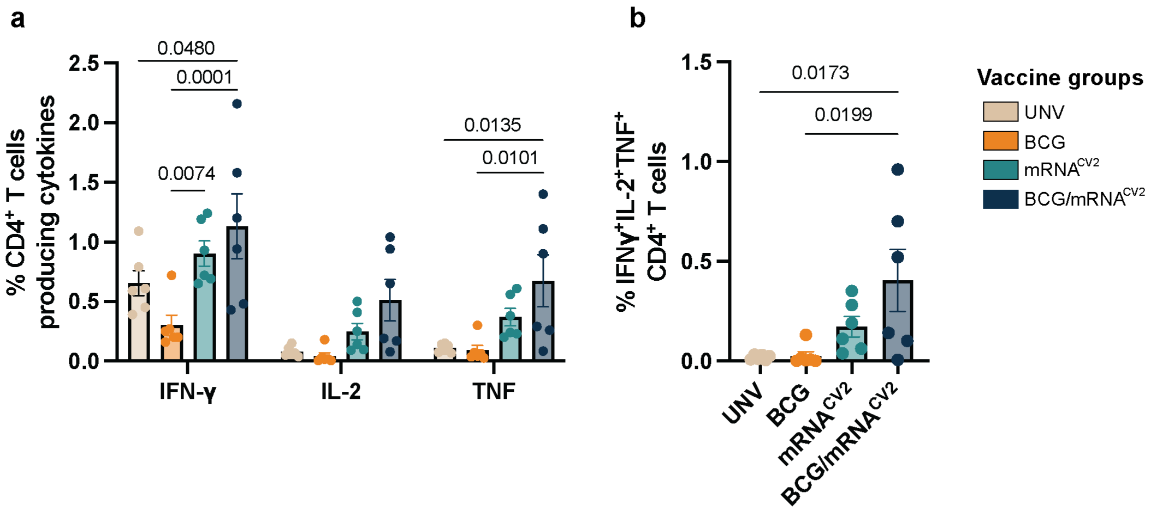


**Supplementary Figure 8: Potent CysVac2-specific CD4$^{+}$ T cell responses of mRNA$^{CV2}$-vaccinated BCG-primed mice in the mediastinal lymph node following *M. tuberculosis* infection.** Proportions of CysVac2-specific **(A)** IFNγ-, IL-2-, or TNF-producing CD4$^{+}$ T cells. **(B)** Proportion of polyfunctional CD4$^{+}$ T cells. Data are expressed as mean +/- SEM with 5-6 mice in each group and representative of 2 independent experiments. Significance determined by **(A)** 2-way ANOVA with Tukey's post-hoc test, and **(B)** 1-way ANOVA with Tukey's post-hoc test.
